# Substantial genetic potential for deep-sea chemoautotrophy extends beyond nitrifiers

**DOI:** 10.64898/2026.01.13.699260

**Authors:** Rebecca S. R. Salcedo, Alexander L. Jaffe, Anne E. Dekas

## Abstract

I.

Microbial chemoautotrophy in the deep sea has the potential to sustain ecosystems at depth, contribute to global carbon sequestration, and ameliorate the currently unbalanced deep-sea carbon budget. However, an understanding of its mechanisms and feasibility is still emerging, particularly beyond that fueled by nitrification. Here, we conduct both a gene-based and genome-resolved analysis of 28 metagenomes in the Northeast Pacific Ocean to investigate the prevalence, distribution, and phylogenetic and metabolic diversity of deep-sea chemoautotrophs. We find that organisms encoding marker genes for dissolved inorganic carbon (DIC) fixation are abundant and widespread at our study site, comprising 11-26% of microbial communities from 150-4000 m water depth. Marker genes for the Calvin cycle and 3-Hydroxypropionate (3HP) bi-cycles are more prevalent than those for the 3-Hydroxypropionate/4-Hydroxybutyrate (3HP/4HB) and reverse Tricarboxylic Acid (rTCA) cycles, the latter two of which are encoded by organisms conducting nitrification. We construct and identify 128 putatively chemoautotrophic metagenome-assembled genomes, spanning 14 phyla including the Proteobacteria, Actinobacteria, SAR324, and the Thermoproteota. They contained genes for the oxidation of carbon monoxide (77.3%), sulfur (68.8%), ammonia (5.5%), and/or methane (3.1%), suggesting diverse catabolisms fuel deep-sea DIC fixation and an underappreciated potential role for aerobic carbon monoxide oxidation. Fifty percent of these genomes encoded multiple inorganic catabolic pathways and 99% included genes for organic matter transport, suggesting catabolic flexibility and potentially facultative autotrophy, respectively. We create an inclusive inventory and map of potential chemoautotrophs at our study site, expanding their known phylogenetic breadth, metabolic repertoires, and potential to impact the carbon cycle.

**Importance:** The deep-sea carbon cycle plays a central role regulating marine ecosystem productivity and the global climate. Chemoautotrophy, the microbial conversion of inorganic carbon (e.g., CO_2_ or HCO^-^_3_) into cellular biomass, is an understudied process with the potential to significantly shift models of the marine carbon budget. While some deep-sea chemoautotrophs are known, a wholistic analysis of organisms with the genetic potential for chemoautotrophy in the dark water column is lacking. Here, we find that microorganisms with the genetic potential for chemoautotrophy are widespread and abundant, and that they are more phylogenetically and metabolically diverse than previously appreciated. In particular, the prevalence of genes involved in CO oxidation suggests a currently unrecognized role in fueling deep-sea chemoautotrophy. By identifying the potential microbial mediators and coupled energy sources, our results allow for more accurate predictions of when, where, and how deep-sea chemoautotrophy occurs, and therefore its potential role in carbon cycling.

## II. Introduction

The biological fixation of inorganic carbon (e.g., CO_2_ or HCO^-^_3_) supports the biosphere and plays a key role in regulating Earth’s climate [1]. While inorganic carbon fixation powered via light energy (photoautotrophy) is most studied and well known, the potential significance of inorganic carbon fixation fueled by the oxidation and reduction of inorganic chemical compounds (chemoautotrophy) is increasingly recognized. Chemoautotrophy in the deep sea may be particularly impactful, playing a role in sequestering atmospheric CO_2_ via the biological carbon pump and providing organic carbon to heterotrophic microbes and macrofauna with limited access to photosynthetically produced organic matter. Additionally, underestimated rates of chemoautotrophy have recently been suggested as one of several potential oversights in the deep-sea carbon budget leading to a perceived imbalance: the known sinks of organic carbon, driven by microbial respiration, currently exceed the known supply from photosynthetic export [2,3].

Chemoautotrophy can be performed by phylogenetically diverse organisms using one of seven biosynthetic pathways for dissolved inorganic carbon (DIC) fixation: i) The Calvin Benson Bassham (CBB) cycle, ii) the 3-hydroxypropionate (3HP) bi-cycle iii) the 3-hydroxypropionate/4-hydroxybutyrate (3HP/4HB) cycle, iv) the dicarboxylate/4-hydroxybutyrate (DC/4HB) cycle, v) the Wood-Ljungdahl (WL) pathway, vi) the reverse tricarboxylic acid (rTCA) cycle, and vii) the reductive glycine (RGP) pathway. Historically, the numerically significant Nitrososphaeria class within the Thermoproteota (formerly Thaumarchaea) have been considered the most significant chemoautotrophs in the deep-water column, pairing autotrophy via the 3HP/4HB cycle to ammonia oxidation in the first step of nitrification [4–6]. However, theoretical estimates of the maximum DIC fixation supported by ammonia oxidation in the Northern Atlantic Ocean are not sufficient to balance the deep carbon budget [2]. Additionally, recent inhibitor experiments demonstrate that ammonia oxidizers fix a minority of total dark DIC fixation in the mesopelagic of the Eastern Tropical Pacific Ocean [7].

Potentially contributing to dark DIC fixation is a growing list of additional deep-sea organisms capable of chemoautotrophy, including Nitrospinae, which perform the second step of nitrification by coupling nitrite oxidation to autotrophy via the rTCA and CBB cycles [8,9], as well as members of the candidate phylum SAR324 and the Gammaproteobacterial order PS1, both of which are thought to couple sulfur oxidation to autotrophy via the CBB cycle [10–14]. Global genomic surveys broadly assessing the metabolic potential of deep-sea microbes have also observed an abundance and expression of genes involved in the CBB, 3HP/4HB, 3HP bi-cycle, and rTCA pathways throughout the water column[12,12,15–17] (Acinas et al. 2021; Lappan et al. 2022; Amano et al. 2024, Jaffe et al., 2025). Despite growing evidence that the genetic potential for chemoautotrophy is not rare in the deep sea, we lack an inclusive inventory of the phylogenetic lineages potentially harboring this ability, what energy sources they harness, and what determines their absolute and relative abundance throughout the water column. Each of the seven pathways for DIC fixation have varying requirements for energy (ATP), substrate (CO_2_ or HCO^-^_3_), and environmental conditions (e.g., oxygen concentrations) (Table 1), raising the possibility that the organisms encoding these different pathways are differentially selected for in the marine environment. However, such trends have not yet been investigated outside of oxyclines (e.g. [19]).

**Table 1:** Characteristics of the seven known carbon fixation pathways[1,19,20,80,112–115]. *Value reported is the same for both pathways, as they are detected as one. Relative abundance reported was determined via gene-based approach, which is unable to distinguish between the DC/4HB and 3HP/4HB pathways.

| DIC Fixation Pathway | ATP Equivalents for Synthesis of Pyruvate (1 Cycle) | Domain | Oxygen Sensitive? | Photo- or Chemo? | DIC Substrate | Detected Here? (% of community) |
| --- | --- | --- | --- | --- | --- | --- |
| Calvin Benson Bassham cycle (CBB) | 7 | Bacteria and Eukaryotes | No | both | CO<sub>2</sub> | 3.2-50.1% |
| Wood Ljungdahl (WL) pathway | 1 | Bacteria and Archaea | yes | chemo | CO<sub>2</sub> | 0.0-0.6% |
| Reverse Tricarboxylic Acid cycle (rTCA) | 2 | Bacteria | yes | both | CO<sub>2</sub> & HCO<sub>3</sub><sup>-</sup> | 0.0-0.6% |
| 3-hydroxypropionate bi-cycle (3HP bi-cycle) | 5-7 | Bacteria | no | both | HCO<sub>3</sub><sup>-</sup> | 3.0-7.4% |
| 3-hydroxypropionate/4-hydroxybutyrate cycle (3HP/4HB) | 6-9 | Archaea | no | chemo | HCO<sub>3</sub><sup>-</sup> | 0.7-6.2%* |
| Dicarboxylate/4-hydroxybutyrate cycle (DC/4HB) | 4-5 | Archaea | yes | chemo | CO<sub>2</sub> & HCO<sub>3</sub><sup>-</sup> | 0.7-6.2%* |
| Reductive Glycine pathway (RGP) | 1-2 | Bacteria and Eukaryotes | No | chemo | CO<sub>2</sub> | not assessed |

We recently reported on the distribution of organisms encoding rubisco (a key enzyme in the CBB cycle) in the deep Northeastern Pacific Ocean [12], but did not investigate the potential for chemoautotrophy outside of the CBB pathway, or the relative abundance of rubisco-encoding organisms compared to those encoding other pathways for inorganic carbon fixation. Here, we investigate the full genetic potential for chemoautotrophy across the natural physical and chemical gradients within the same transect, including 5 vertical profiles (50m-4000m water depth) across 281km off the coast of Northern California. Using 28 deeply sequenced metagenomes (44.3-59 Gbp/sample) we conduct a gene-based analysis to track relative abundance and distribution of genes involved in autotrophic pathways throughout the region, as well as a genome-resolved analysis where we construct genomic bins for putatively chemoautotrophic organisms to investigate their phylogenetic identities and full genetic repertoires. Our study contributes to the growing body of literature on DIC fixation in the deep sea, with the goal of understanding the taxonomic and mechanistic underpinnings of carbon cycling in this largely mysterious majority of the surface of our planet.

## III. Results

### Community Prevalence and Environmental Distribution of Pathways for DIC Fixation

We began by assessing the presence and relative abundance of 25 marker genes (Table S1) associated with six autotrophic pathways (Table 1). These marker genes were identified in assembled contiguous sequences (contigs) from 28 metagenomes collected in the Northeastern Pacific (Figure 1A & B). This gene-based approach maintains all assembled sequences, precluding potential bias incurred when only assessing the subset of reads included in metagenome-assembled genomes (MAGs) (see Figure S1 and discussed below). We did not assess the prevalence of the reductive glycine pathway (RGP), as it currently has no unique marker genes allowing for distinction from its oxidative counterpart [20].

**Figure 1:**
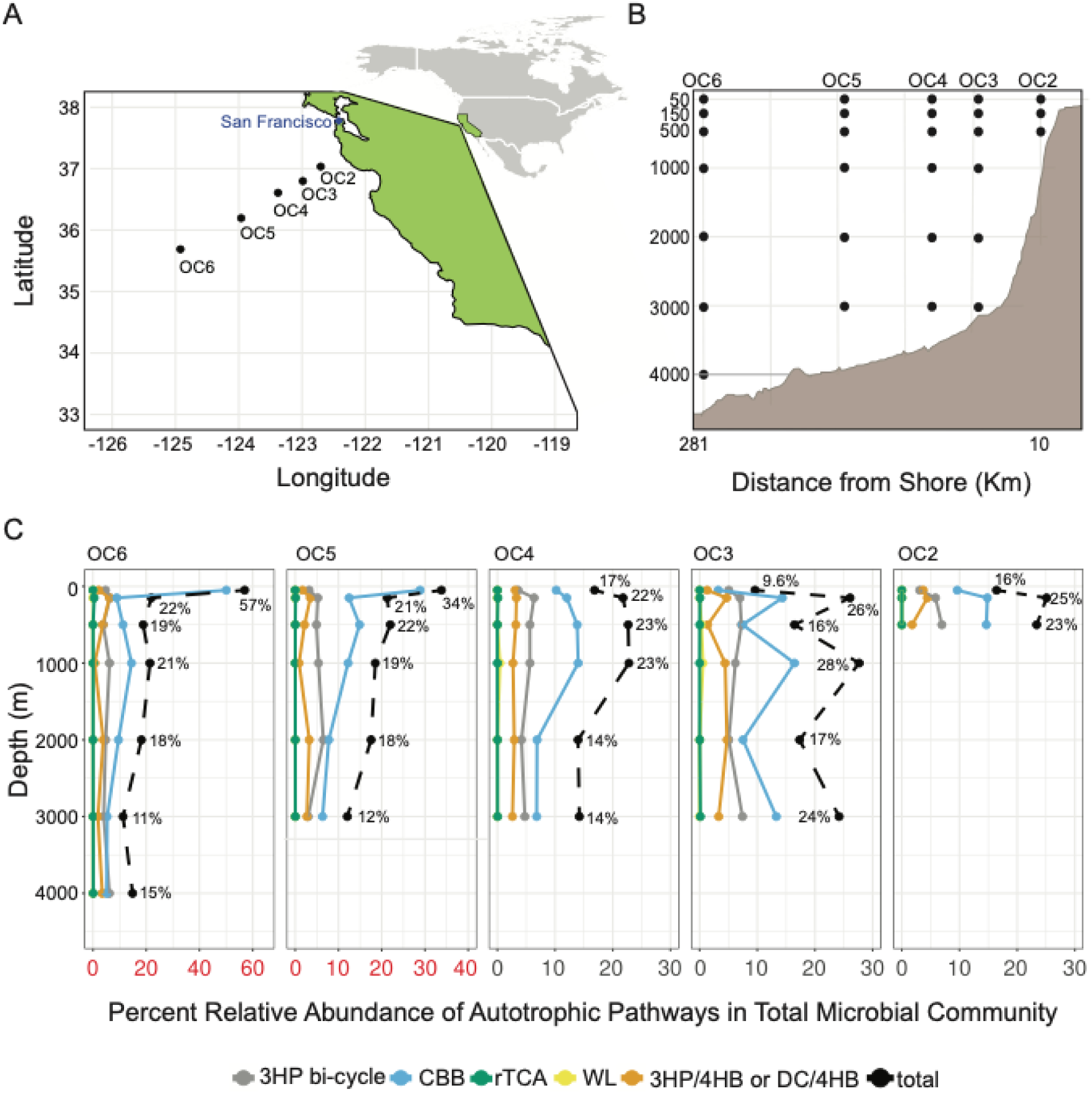
The sampling transect analyzed (A and B), the relative abundance of putative autotrophs as a percentage of the microbial community and the relative abundance of each pathway involved in autotrophy (C). In (A) each black dot represents a metagenome. In (C), red axis indicate axis that have difference scales than the rest of panel C.

We detected genes involved in the CBB cycle, the 3HP bi-cycle, the 3HP/4HB or DC/4HB cycle, the Wood-Ljungdahl (WL) pathway, and the rTCA cycle throughout the transect (Figure 1). Because genetic markers of the DC/4HB cycle overlap with those of the 3HB/4HB cycle, the two pathways are discussed in conjunction in the gene-based analysis. Across all samples, we found that 11-57% (average 21.0%) of microorganisms per sample encoded at least one marker gene for one of the six pathways, determined by normalization of marker gene coverage to the average coverage of four single copy genes (Figure 1C, Table S1). Organisms encoding the CBB cycle, the 3HP bi-cycle, and the 3HP/4HB or DC/4HB cycle were found in 100% of samples investigated and were the most prevalent putative autotrophs (5.1-50.1%, 3.0-7.4%, and 0.7-6.2% of the total communities, respectively) (Table 1). Organisms encoding the rTCA and WL cycles were found in 67% and 14% of samples and had the lowest relative abundances (0.0%-0.5% and 0.0%-0.6% of the total microbial communities, respectively). The CBB pathway was the most abundant autotrophic pathway in all samples except at site OC6, 4000m water depth, where the 3HP bi-cycle reaches 41.5% of the total autotrophic gene relative abundance (6.2% of the total community). Genes involved in the CBB cycle peak in relative abundance in the 50m samples of sites OC5, OC6, and OC4, likely due to an increase in cyanobacteria abundance (Figure S2). Excluding the 50 m samples, organisms encoding the CBB cycle comprise 5.1-16.5% of the communities sampled. Based on gene coverage, the gene-based approach indicates that on average 19.8% of the total microbial community at depths ≥150m (referred to here as dark; <0.045 umol m^-2^s^-1^ photosynthetic active radiation or <0.01% of surface levels) has the genetic potential for autotrophy.

We assessed whether the prevalence of autotrophic potential and/or the relative abundance of specific pathways was correlated with the physiochemical parameters of the water column. These parameters varied substantially across the dataset: hydrostatic pressure (6 – 400 atm), temperature (1.16-14.5 °C), ammonium (not detected to 368 nM), NOx (0-10.5 μM), urea (20.5-1063 nM), and oxygen (0.32-5.87 ml/l) [21]. However, no statistically significant correlations were observed (Table S2, S3). Considering only the dark samples (≥150m), significant negative correlations are observed between the rTCA and NOx at sites OC6 and OC5 (R = -0.97, -0.98, adjusted p-value = 0.010, 0.015 respectively).

### Phylogenetic Diversity and Relative Abundance of MAGs Encoding Genes for DIC Fixation

Next, we generated 742 non-redundant species level MAGs (>50% completeness, <5% contaminated), to link the presence of genes involved in autotrophic pathways to phylogenetic identity and energy-harnessing metabolisms (catabolisms). This MAG set included 7.8-46.1% of the total reads in each metagenome (Figure S1). After manual refinement and curation, we recovered 212 MAGs that contained at least one key gene involved in an autotrophic pathway (see methods), which we refer to as the MAG set encoding marker genes for autotrophic pathways (Table S4). Of those 212, 128 also encoded at least one marker gene for an inorganic chemical catabolic pathway suggesting a chemolithoautotrophic lifestyle (see details of catabolism identification below). We refer to these 128 MAGs as the putatively chemoautotrophic MAG set. Our previous study identified and reported 11 non-redundant rubisco-encoding MAGs from this dataset[12]; here we present an additional 201 MAGs encoding genes involved in three other DIC fixation pathways (3HB bi-cycle, 3HB/4HB, and DC/4HB) (Table S4). Identified MAGs in the putatively chemoautotrophic MAG set span 2 domains (Archaea, 6; and Bacteria, 122) and 14 phyla (Actinobacteriota, 24; Proteobacteria, 68; Chloroflexota, 3; Desulfobacteriota_B, 3; Gemmatimonadota, 2; Hydrogenedentota, 2; JAAXHH01, 1; Myxococcota, 1; Myxococcota_A, 10; SAR324, 6; UBA8248, 2; Thermoproteota, 4; and Halobacteriota, 2) (Table 2). For more than half of the phyla, all MAGs were affiliated with a single family, including the Thermoproteota (Nitrosopumilaceae), SAR324 (NAC60-12), Gemmatimonadota (UBA6960), Halobacteriota (UBA12382), Hydrogenedentota (GCA−2746185), JAAXHH01 (JAAXHH01), and Myxococcota (JABJBS01), while others contained more, and up to 24 in the case of the Proteobacteria (Table 2, Figure S3).

**Table 2:** Description of the putatively chemoautotrophic MAG set, i.e. MAGs containing a marker gene for both an inorganic carbon fixation pathway and an inorganic catabolism. *with the exception of the phylum Proteobacteria, which is displayed at class level.

| Domain | Phylum* (families, n=) | MAGs (n=) | Pathway(s) Detected | Mean Relative Abundance (%) | Maximum Relative Abundance (%) |
| --- | --- | --- | --- | --- | --- |
| Bacteria | Actinobacteriota (4) | 24 | CBB, 3HP Bicycle | 4.50 | 10.01 |
| Bacteria | Alphaproteobacteria (13) | 35 | CBB, 3HP Bicycle | 1.45 | 5.53 |
| Bacteria | Chloroflexota (2) | 3 | CBB, 3HP Bicycle | 0.11 | 0.67 |
| Bacteria | Desulfobacterota_B (1) | 3 | 3HP Bicycle | 0.09 | 0.18 |
| Bacteria | Gammaproteobacteria (11) | 33 | CBB, 3HP Bicycle | 3.34 | 6.89 |
| Bacteria | Gemmatimonadota (1) | 2 | 3HP/4HB or DC/4HB | 0.26 | 0.69 |
| Archaea | Halobacteriota (1) | 2 | DC/4HB | 0.04 | 0.13 |
| Bacteria | Hydrogenedentota (1) | 2 | 3HP Bicycle | 0.07 | 0.21 |
| Bacteria | JAAXHH01 (1) | 1 | 3HP/4HB or DC/4HB | 0.08 | 0.21 |
| Bacteria | Myxococcota (4) | 1 | CBB, 3HP Bicycle, 3HP/4HB or DC/4HB | 0.06 | 0.22 |
| Bacteria | Myxococcota_A (1) | 10 | 3HP/4HB or DC/4HB | 0.16 | 0.25 |
| Bacteria | SAR324 (1) | 6 | CBB | 3.19 | 5.02 |
| Archaea | Thermoproteota (1) | 4 | 3HP/4HB | 2.32 | 4.39 |
| Bacteria | UBA8248 (2) | 2 | 3HP/4HB or DC/4HB | 0.06 | 0.21 |

Unlike in the gene-based analysis, we could differentiate between the 3HP/4HB and the DC/4HB cycles here using genomic context: four contigs with genes for the 3HP/4HB or DC/4HB cycle binned into MAGs classified as Thermoproteota, a lineage well documented to encode a modified 3HP/4HB cycle [22]. On the other hand, fifteen contigs associated with the 3HP/4HB or DC/4HB consistently affiliated with NCBI phylum Euryarcheota (aka Halobacteriota) (Figure S2, Table S5), whose representatives have been found with the DC/4HB in aquifers [23]. We assessed the degree of completion of each carbon fixation pathway in our 2 Halobacteriota MAGs, which revealed a more complete DC/4HB than 3HP/4HB cycle (Figure S4). Similarly, we observe a more complete DC/4HB pathway than 3HP/4HB in our one Myxococcota MAG. Thus, we infer our Halobacteriota MAGs, all from family UBA12382, and our Myxococcota MAG encode the DC/4HB cycle while our Nitrosophaeria MAGs encode the 3HP/4HB cycle.

We observe taxa-specific relative abundance depth trends within the putatively chemoautotrophic MAGs throughout the water column. The biggest changes were observed between 50 and 150 m water depth, above and below the photocline, but trends at both phyla and family levels were evident throughout the dark ocean as well. Putative chemoautotrophic Actinobacteriota peak in relative abundance at 500m depth across all sites (0.3-9.5%), driven by the distribution of the MedAcidi−G1 family, and reach the highest relative abundance of any putative chemoautotrophic phylum across the dataset at 10% at OC3:500m (Figure 2A, S3).

**Figure 2:**
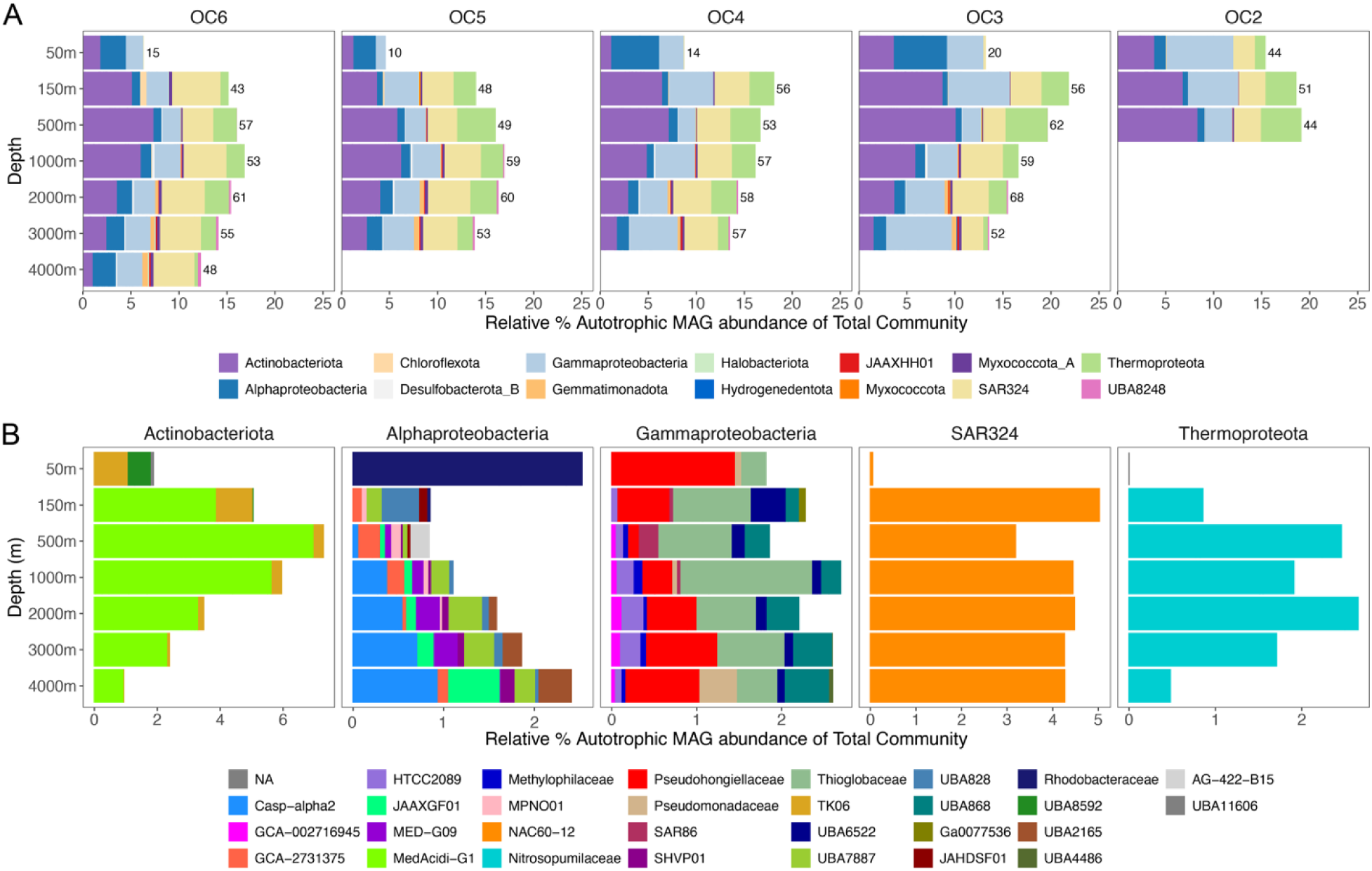
A) Percent relative abundance of putatively chemoautotrophic MAGs across the transect relative to the total community, grouped by taxonomic phyla, except for the Proteobacteria, which is listed at the class level. Numbers indicate number of MAGs detected in sample. B) Percent relative abundance of putatively chemoautotrophic MAGs (relative to total community) across site OC6 of the 5 most abundant taxonomic groups, color coded at the family level (unless family was not identified, then it is colored by order level).

SAR324 comprises a stable 2.3-5.0% of the microbial community throughout samples 150 m water depth and deeper (Figure 2A). Putative chemoautotrophic Alphaproteobacteria peak in relative abundance in the 50m samples (5.5% at OC3:50m), then decrease with depth before increasing again to 2.4% in the bathypelagic (OC6:4000m). Interestingly, while most Alphaproteobacterial families increase in relative abundance with depth, a single family, the Rhodobacteraceae, is very abundant at 50m (5.4% at OC3:50m) and decreases with depth, resulting in a C-shape abundance pattern at the phylum level. Putative chemoautotrophic Gammaproteobacteria are dominated by the family Thioglobaceae (within order PSI) and range from 0.5-4.1% relative abundance of the total microbial community below 50m. Putative chemoautotrophic Nitrosopumilaceae (within the Thermoproteota) MAGs peak in relative abundance at 500 m water depth across sites (with the exception of Site OC6) and reach a maximum of 4.5% at OC3:500m. Several taxa peak in relative abundance in the deepest samples, including putative chemoautotrophic MAGs within the Gemmatimonadotal family UBA6960 (0.7% at OC6:4000m) and putative chemoautotrophic MAGs within the family Hydrogenedentota GCA−2746185 (0.2% at OC3:3000m) (Figure S3).

### Catabolic Pathways Within Putatively Chemoautotrophic MAGs

To examine energy sources that could power DIC fixation, we investigated the presence of a large set of marker genes associated with the oxidation of inorganic compounds (Table S6) across the set of 212 MAGs encoding a marker gene for a DIC fixation pathway. These include but not limited to genes encoding the potential for oxidation of ammonia, nitrite, multiple forms of reduced sulfur (thiosulfate, sulfide, sulfite, and elemental sulfur), carbon monoxide (CO), methane, and hydrogen (Figure 4, Table S6). Based on the recovery of at least one marker gene for a given catabolism, we identify 128 MAGs that encode both an DIC fixation pathway and an inorganic catabolism (i.e., the putatively chemoautotrophic MAG set). We observe 77.3% (99/128) of putatively chemoautotrophic MAGs contained potential for aerobic CO oxidation(*coxMSL*), 5.5% (7/128) for ammonia oxidation (*hao/amo*), 68.8% (88/128) for one of various sulfur oxidation pathways(*apr/sat, sox, fccB/sqr, or dsrAB*), and 3.1% (4/128) for methane oxidation (Figure 3AB). Potential for sulfite oxidation was associated only with the 18 MAGs (14.1%) from Alpha- and Gamma-proteobacteria, SAR324, and UBA824 that encoded both *aprA* and *sat*, as *sat* alone is also used in assimilatory sulfate metabolism [24] (Figure S5). We did not detect genes for the oxidation of nitrite and hydrogen in our putatively chemoautotrophic MAGs.

**Figure 3:**
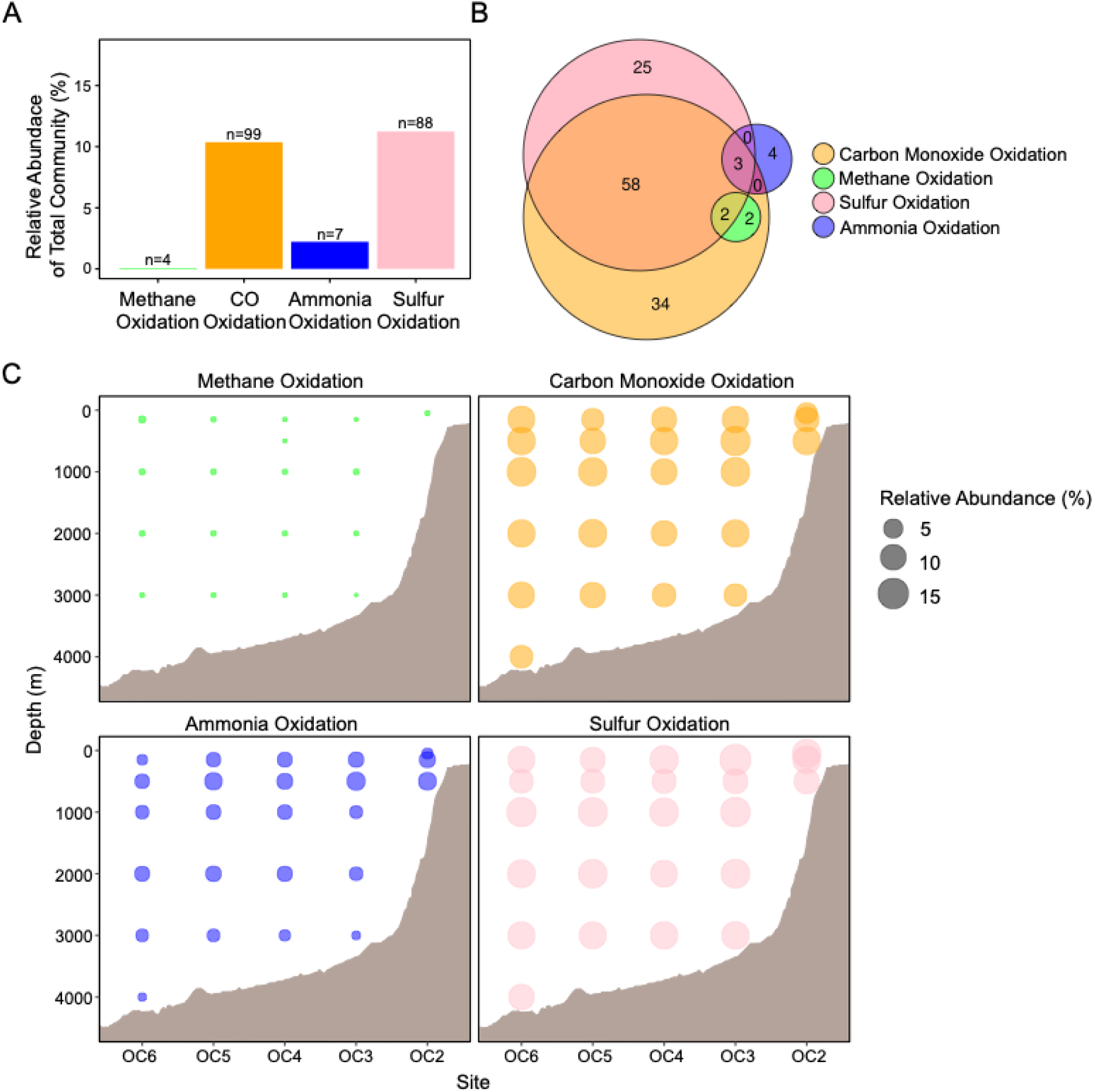
Relative abundance and distribution of catabolic pathways throughout the transect. (A) Relative abundance of putatively chemoautotrophic MAGs with one of four catabolisms (n=128 genomes) (B) Venn diagram displaying number of putatively chemoautotrophic MAGs containing any given catabolic pathway or combination of pathways. (C) Spatial distribution of the prevalence of putatively chemoautotrophic MAGs with various catabolisms. Ocean floor is approximate, estimated with GeoMapApp[116].

**Figure 4:**
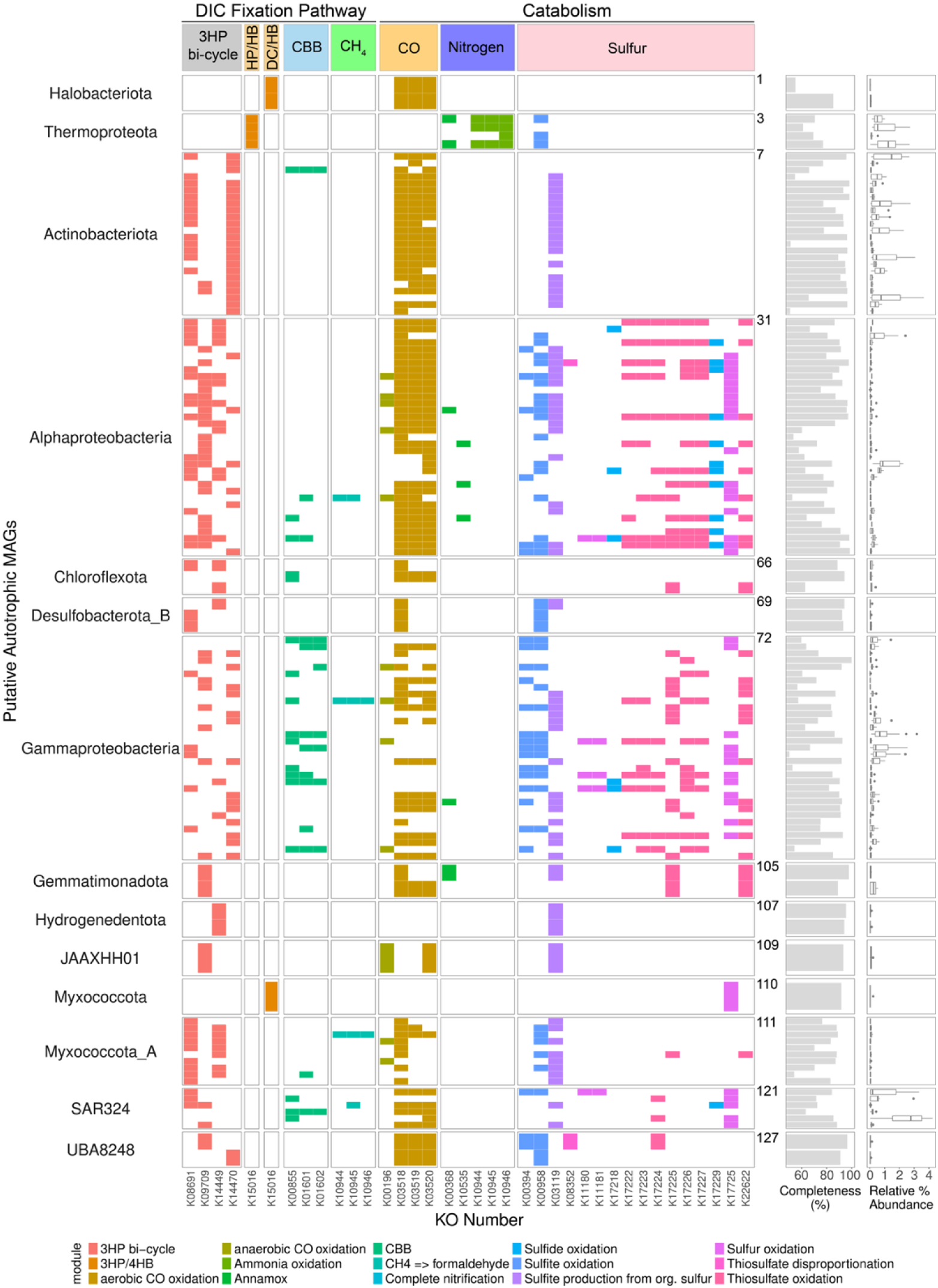
Metabolic potential of putatively autotrophic MAG set that encodes both a pathway for inorganic carbon fixation and an inorganic catabolism. Each row represents a single MAG, grouped by taxonomic lineage. Each column represents a single KO associated with a given catabolism or pathway of interest, grouped into larger metabolic functions. Boxes are colored coded by KEGG module aka metabolic pathway. MAG completeness and associated range of percent relative abundance across the transect are displayed to the right. 30 KOs identified in a total of 128 MAGs. KO K15016 is listed twice, as it can play a role in both the 3HP/4HB and DC/4HB pathways. Number to the right of presence/absence plot corresponds to “short_id” MAG label in supplementary table S4; the first MAG in each taxonomic lineage is labeled. Acronyms: DIC = Dissolved Inorganic Carbon.

Interestingly, 50.7% of putatively chemoautotrophic MAGs with a recovered catabolism encoded multiple catabolic pathways, with CO and sulfur oxidation being the most common pairing (45.2%) (Figure 3B). Putatively chemoautotrophic MAGs with methane monooxygenase were only found with either both aerobic CO dehydrogenase (CODH) and sulfur oxidation genes or just CODH, while genes involved in ammonia oxidation were often found with no other supporting catabolism, as expected of the ammonia-oxidizing Nitrosopumilaceae [25]. Three MAGs (2.3%) house genes for three identified catabolisms: CO oxidation (*coxMSL*), nitrogen oxidation (*hao*), and sulfur oxidation (*sox*) (Figure 3B, 4). All 3 fall within the Alphaproteobacterial family UBA7887, a newly discovered and poorly-understood lineage of marine Alphaproteobacteria [26,27].

When assessing the relative abundance of putatively chemoautotrophic MAGs as a function of their paired catabolism, we observed a dominance of CO and sulfur oxidation across our transect (Figure 3A, C). Depending on sample, putatively chemoautotrophic MAGs encoding CO oxidation genes account for 6.3-13.5% of the total microbial community, and those with sulfur oxidizing genes account for 8.4-15.4% (Figure 3C). The distribution of putatively chemoautotrophic MAGs encoding *coxSML* is relatively consistent with depth and distance from shore, with its highest relative abundance in the mesopelagic (500m-1000m). Relative abundance of putatively chemoautotrophic MAGs capable of sulfur, sulfite, or thiosulfate oxidation consistently drop in abundance at 500m water depth before increasing again at 1000 m. Putatively chemoautotrophic MAGs with the potential ability for ammonia oxidation account for 0.5-4.5% of the total microbial community (Figure 3C), with the highest relative abundance near 500m.

### Prevalence of Genes Encoding ABC Transporters in the Putatively Chemoautotrophic MAG Set

To assess the potential for transport and utilization of externally derived organic matter in our putatively chemoautotrophic MAG set, we analyzed the presence of ATP-binding cassette (ABC) transporters with known organic matter substrates (Table S6, Figure S5). We see that 127/128 of MAGs encode at least one organic matter associated ABC transporter, with transport of branch chained amino acids (*hmgL*/K01640, *ilvE*/K00826, and *pdhD*/K00382) being the most prevalent (122/128). Monosaccharide and oligosaccharide associated ABC transporters are also widespread across our putative chemoautotrophic MAG set (40/128), especially among the Proteobacteria, Hydrogenedota, SAR324, JAAXHH01, and UBA8248. We did not recover ABC transporters in only one MAG, which belonged to Actinobacteriota, and was only 65.7% complete. In addition to ABC transporters, 120/128 putative chemoautotrophic MAGs encode genes for the glycine cleavage system (Figure S5).

## IV. Discussion

The deep-sea carbon cycle is foundational to our planet’s chemistry and climate, but it remains understudied relative to the more productive and easily accessible surface ocean. A discrepancy in our understanding of the deep-sea carbon cycle has been increasingly recognized: the known sinks of organic carbon, driven by microbial respiration, exceed the known supply from photosynthetic export and inferred rates of *in situ* production (nitrification-based chemoautotrophy) [2,3]. This discrepancy may be explained by multiple oversights in our conceptual understanding and quantitative accounting, including the potential for more diverse, and/or active chemoautotrophs in the dark ocean [3,28]. The implications of substantial production of organic matter at depth are significant for both marine ecosystems and global biogeochemistry, particularly if it is energetically supported by diverse, widely available chemical energy sources that allow it to be robust temporally and spatially. Advancing such an understanding of deep-sea chemoautotrophic physiology with genomics thereby facilitates more accurate models of the carbon cycle and predictions of how dark carbon fixation will respond to future natural and anthropogenetic changes. Specifically, establishing the abundance, distribution, and phylogenetic and metabolic diversity of potential chemoautotrophs in the deep sea with genomics can guide future efforts to obtain fixation rates and contextualize those rates with the physiological mechanisms underlying them.

Here, we use complementary gene- and genome-based approaches to show widespread autotrophic potential throughout the water column off the coast of Northern California. Our gene-based approach provides an inclusive overview of this potential by not restricting the analysis to reads represented by the MAG set (7.8-46.1%), and our genome-based approach provides genomic context to pair potential with taxonomic identity and supporting catabolism. In both approaches we were inclusive by not requiring pathway completeness. Although this approach increases the chance for false positives, it avoids significant false negatives arising from genome incompleteness and the misannotation of genes in divergent chemoautotrophs expected in the genetically novel deep sea microbiome [15]. The potential for poor pathway recovery due to misannotation is particularly high for pathways involving 3-hydroxypropionate intermediates, which are prone to incomplete recovery due to known logistical challenges with protein annotation (e.g., [29]) and are often fragmented in marine datasets [30]. Consistent with this, 5/6 of the marker genes for the 3HP/4HB pathway in the 4 Nitrosopumilaceae MAGs recovered here were not recovered, while there is extensive empirical evidence of a functional 3HP/4HB cycle in this lineage. Inferences of autotrophy based on the detection of marker genes rather than complete pathways are common in environmental samples (e.g., [12,31], but detection of certain marker genes (e.g. rubisco forms I/II for CBB, *aclAB* for rTCA) are likely more indicative than others. Increased knowledge of the biochemistry and sequence families of the enzymes in these pathways, particularly the lesser studied ones, will likely improve inferences based on marker gene recovery.

To strengthen our predictions of chemoautotrophy while maintaining a wide net, we required genomic evidence (detection of a marker gene) for both an inorganic chemical catabolism and a DIC fixation pathway for a MAG to be classified as putatively chemoautotrophic. Our list of assessed catabolisms is not exhaustive and could therefore exclude bona-fide chemoautotrophs, but we have additional confidence in chemoautotrophic potential of the retained MAGs. This step reduced our set of 212 MAGs containing a marker gene for DIC fixation to 128 putatively chemoautotrophic MAGs, and excluded, for instance, all 57 Thermoplasmatota (MGII) MAGs in which we detected a marker gene for the 3HP/4HB or DC/4HB pathway (Table S4). Previous studies have also recovered marker genes for autotrophy within the Thermoplasmatota, including evidence for the WL, rTCA, and 3HP/4HB or DC/4HB pathways [32–36], but they are widely considered heterotrophic due to presence of organic matter processing genes and association with particles [37,38]. Ultimately, physiological tests are necessary to confirm the metabolic lifestyle and capabilities of microbial lineages, either in culture or through manipulation of environmental samples (e.g. [39,40]). The work presented here identifies, quantifies, and characterizes potential chemoautotrophs, highlighting their potential significance and guiding physiological and rate-based future studies.

### An Abundant, Phylogenetically Diverse Microbial Community of Putative Marine Chemoautotrophs

The putatively chemoautotrophic MAG set is comprised of a diverse group of organisms spanning 14 phyla, including known chemoautotrophs such as the Thermoproteota (using the 3HP/4HB), SAR324 (using the CBB), and Gammaproteobacterial order PSI (using the CBB) [12–14,41–43]. We observe a dominance of the CBB pathway over other pathways for DIC fixation, followed by the 3HP bicycle, throughout the vertical water column in both the gene- and genome-resolved approaches. This is contrary to the notion that nitrifying organisms (encoding the 3HP/4HP or rTCA cycles) are the most abundant chemoautotrophs at depth, but is consistent with a recent study indicating that nitrifiers contribute a minority of total inorganic carbon fixation in the mesopelagic [7]. Previous molecular studies finding high relative abundances of CBB-encoding organisms at depth [11–13,15]—with additional evidence displaying active transcription of rubisco [11,12] or direct DIC assimilation [13]—further highlight the potential importance of CBB-driven DIC fixation in the deep sea.

Our study also suggests an expanded phylogenetic distribution of the 3HP bi-cycle and DC/4HB pathways. We identify 2 putatively chemoautotrophic MAGs from the Hydrogenedentota phyla (formerly Hydrogenedentes) encoding marker genes for the 3HP bi-cycle. This pathway was thought to be phylogenetically restricted to the *Chloroflexi*, but has now been found across multiple phyla [32]. The Hydrogenedentes are a newly defined phylum, being found to encode the rTCA, the CBB, and the WL pathway [44,45], and now we observe the 3HP bicycle (Figure 4). Similarly, the DC/4HB cycle was thought to only be found in archaea [1], yet we recover K15016, a marker gene for the DC/4HB and 3HP/4HB pathways, within bacterial lineage Myxococcota (Figure 4). Our data are consistent with a new but growing body of literature identifying this pathway in the bacterial domain [32,46–48] and the first documented evidence for the DC/4HB cycle in the Myxococcota phyla to our knowledge.

### Catabolisms Potentially Fueling Chemoautotrophy

Consistent with the low relative abundance of genes in the 3HP/4HP and rTCA cycles, we observed low relative abundance of lineages capable of nitrification in the putative chemoautotrophic MAG set. We report a relative abundance of 0.5-4.5% of MAGs encoding genes for ammonia oxidation, and a relative abundance max of 0.019% of contigs classified as NCBI phylum Nitrososphaerota, both noticeably lower than values previously reported at Station ALOHA, Pacific Ocean, for AOA (up to 39%; [49]), and even a previous 16S rRNA-based survey at our study site (∼20% Thermoproteota/Crenarchaeota; [21]). Genetic micro-diversity negatively impacts marine AOA assembly and binning [50] suggesting relative abundance values for AOA based on metagenomics, such as ours, may be underestimates. We also detected no high-quality MAGs of nitrite-oxidizing bacteria (NOB) within our putatively chemoautotrophic MAG set, consistent with a low abundance of unbinned contigs associated with NOB (0.13-0.21% of the autotrophic community). Unlike for AOA, 16S rRNA gene-based abundance assessments of NOB at this site were also low [21]. However, NOB have increased enzymatic efficiency and large cell size, allowing them to perform impactful geochemical processes while maintaining low numerical abundance in the microbial community [8]. Together, these data indicate that while nitrifying organisms are certainly a component of chemolithoautotrophic metabolism at this site, they comprise a minority of the diversity and likely a minority of the total chemoautotrophic cells.

We observed extensive co-occurrence of genes involved in DIC fixation and the oxidation of various reduced sulfur compounds. Oxidation of reduced sulfur compounds have been predicted to power DIC fixation in organisms encoding the CBB previously [11–15] and DIC fixation has been stimulated with thiosulfate additions in seawater [51]. Here we show widespread genes for sulfur oxidation in MAGs encoding DIC fixation pathways other than the CBB, expanding both the known metabolic combinations and the phylogenetic breadth of these organisms in the ocean. In particular, we see 78/128 MAGs encoding genes for both sulfur oxidation and the 3HP bi-cycle (Figure 4). Though oxidation of sulfur has not been directly linked to DIC fixation via the 3HP bi-cycle in cultured isolates, a previous genomic survey has observed MAGs co-encoding these two pathways across multiple non-marine habitats [32], and a metatranscriptomic investigation of *Chloroflexota* populations shows co-expression of sulfide oxidation and 3HP-bicycle genes in hot springs [52]. Our study highlights the potential importance of sulfur oxidation in supporting DIC fixation in the deep sea, and particularly beyond organisms encoding the CBB cycle.

We observe the *coxMSL* gene cluster, which encodes three subunits of the aerobic CODH, to be widely distributed across both the phylogenetic breadth of our putatively chemoautotrophic MAG set (Figure 4) and the physical environment (Figure 3C), suggesting a currently underappreciated role for CO oxidation in marine chemoautotrophy. Widespread distribution of the *coxMSL* gene cluster in the water column has been previously reported; found down to 4000m at Station ALOHA [53], across the global Malaspina sampling effort [15], the deep Mediterranean sea [54–56], and hadal trenches [57], but is typically interpreted as an energy supplement for heterotrophs [58–60]. However, the detection of *coxMSL* in putatively chemoautotrophic MAGs suggest it could support chemoautotrophy in the marine water column as well. It is sufficiently energetically favorable to support autotrophy, generating 248.7 kJ/mol CO oxidized [61]. This is a similar energetic yield to the that of the aerobic oxidation of hydrogen, which is shown to fuel marine chemoautotrophic and mixotrophic metabolism [16,62]. Our detection of *coxMSL* genes in 99 (more than 75%) of our putatively chemoautotrophic MAG set proposes that CO oxidation could be an underappreciated fuel for chemoautotrophy in the dark ocean, either as a sole or contributing catabolism. Notably, of the 99 non-redundant MAGs, 34.3% of those have no other encoded catabolism detected, even among those with completeness values of ≥95%, suggesting it may be the sole fuel for DIC fixation in these organisms. While not yet demonstrated in a marine water column isolate, chemoautotrophic growth supported solely by aerobic CO oxidation is common in other environments including soils [61], hot springs [63], alkaline lakes [64], deep-sea caldera sediment [65], and freshwater systems [66]. Additionally, a recent incubation-based study displayed bulk CO oxidation in seawater samples from 1 m water depth, as well as co-expression of *coxL* and rubisco across the TARA sampling effort *[18]*. Further research is required to determine whether marine chemoautotrophs are capable of harnessing energy from CO and if environmental sources of it are large enough to support them in the deep. Sources of marine CO include photolysis in the surface ocean [67], hydrothermal venting and seafloor spreading at the seafloor [58], and direct dark microbial production through currently unknown mechanisms [67], but concentrations of CO throughout the water column are not currently known.

### Evidence for Catabolic Flexibility and Organic Matter Use in Deep-sea Chemoautotrophs

The widespread detection of genes involved in organic matter transport within the putatively chemoautotrophic MAG set indicates extensive use of organic matter and potentially auxotrophy and/or facultative chemoautotrophy. Uptake and use of amino acids is common in phototrophs [68], and is known to occur within chemoautotrophs, as well (e.g., AOA [69,70]). Genes for organic matter transport in chemoautotrophs can but does not necessarily indicate auxotrophy, the inability to produce certain biomolecules that therefore must be sourced externally, or facultative chemoautotrophy, the ability to switch between primarily organic and inorganic metabolism. The potential for facultative chemoautotrophy in deep-sea organisms has been suggested previously based on co-encoding of genes involved in chemoautotrophic and heterotrophic pathways and with non-constitutive rubisco expression [12,15,42]. The near ubiquitous presence of amino acid-specific ATP-binding cassette (ABC) transporters indicates the potential importance of external amino acids for marine chemoautotrophs, and the presence of mono- and oligosaccharide-specific ATP-binding cassette (ABC) transporters in about a third of the putative chemoautotrophic MAG set further indicates the importance of additional types of organic matter in a subset of these organisms. These findings highlight to potential for facultative chemoautotrophy, but transcriptional and/or physiological studies are necessary to establish how and when these organic substrates are used. Genes for the glycine cleavage pathway were also nearly ubiquitous (Figure S5), and suggests a widespread ability of deep-sea chemoautotrophs to supplement inorganic metabolism with energy gleaned from glycine cleavage, a mechanism previously proposed in heterotrophs as an adaptive metabolic strategy for the oligotrophic deep sea [15,71,72].

Expanding beyond facultative or auxotrophic chemoautotrophy, our data provide evidence for the lesser explored catabolic flexibility within chemoautotrophs, or the ability to harness multiple inorganic energy sources to support DIC fixation. We found that 50.7% of putative chemoautotrophic MAGs possess marker genes for multiple inorganic catabolisms (Figure 3B), suggesting that catabolic flexibility is common within deep-sea chemoautotrophs. This metabolic flexibility could provide unique advantages in low-nutrient systems or environments with a high degree of nutrient flux [1,15].

Harnessing two inclusive metagenomic analyses we reveal a phylogenetically expansive community of organisms potentially capable of chemoautotrophy fueled by a variety of sources of chemical energy. Activity assays and isolation attempts targeting these groups is now needed to probe further into their metabolic lifestyle and their potential contribution to global carbon cycles.

## V. Materials and Methods

### Sample collection, DNA Extraction and Sequencing

Methods for seawater sampling, DNA extraction, and metagenomic sequencing were previously described in [73] and[12]. Briefly, sweater was collected aboard *R/V Oceanus* in May 2017, filtered through 0.2μm Sterivex filters, flash frozen in liquid nitrogen and stored at -80C before extraction and sequencing. Raw reads were accessed through NCBI bioproject: PRJNA1054206.

### Relative abundance and distribution of genetic potential

Sequences were quality controlled with FastQC [74] and bbduk.sh v39.01 [75], assembled with MEGAHIT v1.2.9 [76], then annotated with prodigal v2.6.3 [77] and kofamscan v1.3.0 [78,79]. Marker genes involved in autotrophic pathways were identified with metabolic reconstruction programs and literature review [1,15,19,80–83]. Annotations of DIC fixation pathways marker genes were curated by requiring ≥80% of default bit score threshold and ≥70% average gene length. Any K01601 sequences identified as eukaryotic (see contig based taxonomy methods section) were removed from downstream coverage calculations. Gene-wise coverage was calculated by inStrain v1.7.1 [84], and marker gene relative abundance was calculated by dividing their coverage by the average coverage of 4 single copy genes: *recA, secY, rpsC, pheS* [85]. DIC fixation pathway relative abundance was determined by averaging each DIC fixation pathways’ marker gene relative abundance. The total DIC fixation relative abundance was calculated by summing all pathways’ relative abundances together.

Correlations between pathway specific/total relative abundance and geochemical parameters [21] were determined via calculation of Pearson’s R Correlation Coefficient with Bonferroni’s correction in R [86,87].

### Generation of MAGs and Assessment of Paired Catabolisms

Metagenome-assembled genomes (MAGs) were constructed as described in Jaffe et al., 2025. Non-redundant MAGs with ≥50% completeness, and ≤5% contamination, as determined by CheckM v1.2.2 [88], were then annotated using prodigal v2.6.3 and kofamscan v1.3.0 and taxonomically classified with GTDB-Tk v2.3.2 [89] before being screened for marker genes involved in multiple metabolisms including but not limited to autotrophy, nitrogen cycling, and sulfur cycling (Table S6). ABC transporters with known organic matter substrates were identified from [90]. Annotation of any gene required bit-score ≥ default KEGG threshold score, and annotation of genes involved in autotrophy also required a gene length ≥70% average gene length. For autotrophic genes, we used custom thresholds (Table S1). MAGs with ≥50% completeness, ≤5% contamination, and at least one DIC fixation marker gene were selected for downstream analysis and manually curated using anvi-refine within Anvi’o v7 [91]. Mean sequencing coverage of refined MAGs was calculated with CoverM v0.6.1 [92], requiring a coverage breadth ≥50% and read ANI of ≥95%.

### Curation of Genes Involved in Nitrogen, Sulfur and Hydrogen Catabolisms

Putative ammonia monooxygenase and methane monooxygenase (amoA/pmoA) protein sequences were distinguished by aligning to a reference set of copper monooxygenases [93] with MUSCLE v5.1 [94], before a tree was built and visualized in Geneious Prime v2023.1.2 [95]. Putative nitrite oxidoreductase and nitrate reductases (*narH*/*nxrB*) were aligned to a reference set of the DMSO superfamily [96] and placed on a tree to identify reaction directionality. Putative *dsr*AB genes were assessed for their oxidative potential through comparison to a comprehensive database of functionally annotated prokaryotic gene clusters[97], as described in [12]. Putative sulfur dioxygenase genes were validated through identification of key functional residues, as described in [12]. Lastly, we searched MAGs for energy-yielding hydrogenases using the reference set and annotation protocols described in [16]. A MAGs was required to encode both a DIC fixation pathway and an inorganic catabolism to be classified as putative chemoautotrophic. Results of genome annotation presented in Table S7.

### Contig-based Taxonomy

Contig taxonomy was inferred using a published approach that assigns putative taxonomic affiliation using the consensus of individual proteins[12,73]. Briefly, predicted proteins on each contig were compared to UniRef100 [98] using DIAMOND v2.0.13.151 [99]. Resulting alignments were filtered (e-values ≤1e-20 and ≥70% coverage) and the most common phylum and class level affiliations were computed for each contig. Manual curation was performed for short contigs or those with ambiguous taxonomic signals and was aided by clustering of contigs by sequence similarity using dRep v3.4.0 (*-pa 0.80 -sa 0.95) [100]*. Only those contigs ≥2500 bp were retained for downstream analysis. Contig mean coverage and coverage breadth was calculated with inStrain profile function[84]. Results are presented in Table S5.

All plots were generated in R with the ggplot2 package [86,101,102]. Other R packages utilized include tidyverse[102], data.table[103], ggthemes [104], patchwork [105], RColorBrewer[106], Biostrings[107], ggh4x[108], VennDiagram[109], eulerr[110], and pals [111]. All code for data analysis is available here (https://github.com/rebeccasophiasalcedo/oc1703wc_deepauto, https://github.com/alexanderjaffe/urea-genomics, and https://github.com/alexanderjaffe/rubisco-genomics).

## Data Availability Statement

Newly constructed putatively chemoautotrophic MAGs and trimmed metagenomic reads are available at NCBI at bioproject: PRJNA1054206.

## Acknowledgements

We thank Doug Bartlett, Francisco Rodriguez-Valera, and past and current members of the Dekas lab for helpful conversations. We also thank the captain and crew of the R/V *Oceanus* for assistance with sample collection. We acknowledge the Stanford Computing Center and the Stanford Shared Geomicrobiology Facility (RRID:SCR_025000) for respective informatic and wet lab support. RSRS was supported by a Stanford Graduate Fellowship. Sampling and sequencing efforts were supported by National Science Foundation awards OCE-1634297 and OCE-2143035 (both to A.E.D). A.L.J. was funded by the Stanford Science Fellows program and the NSF Postdoctoral Fellowship in Ocean Sciences. A.E.D. was supported by NSF award OCE-2143035.

## Supplemental material

Table S1: list of key genes and KO associated with autotrophic pathways

Table S2: Pearson R and adjusted p-values for correlation analysis of total gene abundance

Table S3: Pearson R and adjusted p-values for correlation analysis of total gene abundance

Table S4: summary of autotrophic MAGs generated

Table S5: list of KO’s screened for across putative chemoautotrophic MAGs

Table S5: genes involved in autotrophy on contigs in dataset

Table S6: list of KO’s screened for across putative chemoautotrophic MAGs

Table S7: KO’s from table S4 found in putative chemoautotrophic MAGs

Figure S1: percentage of reads mapping to assembly vs bin set

Figure S2: Presence and taxonomic identity of scaffolds encoding genes involved in autotrophic pathways across the transect

Figure S3: relative abundance of putative chemoautotrophic MAGs broken down by family

Figure S4: complete C fixation pathways

Figure S5: sulfite oxidation genes

**Supplementary Figure S1:**
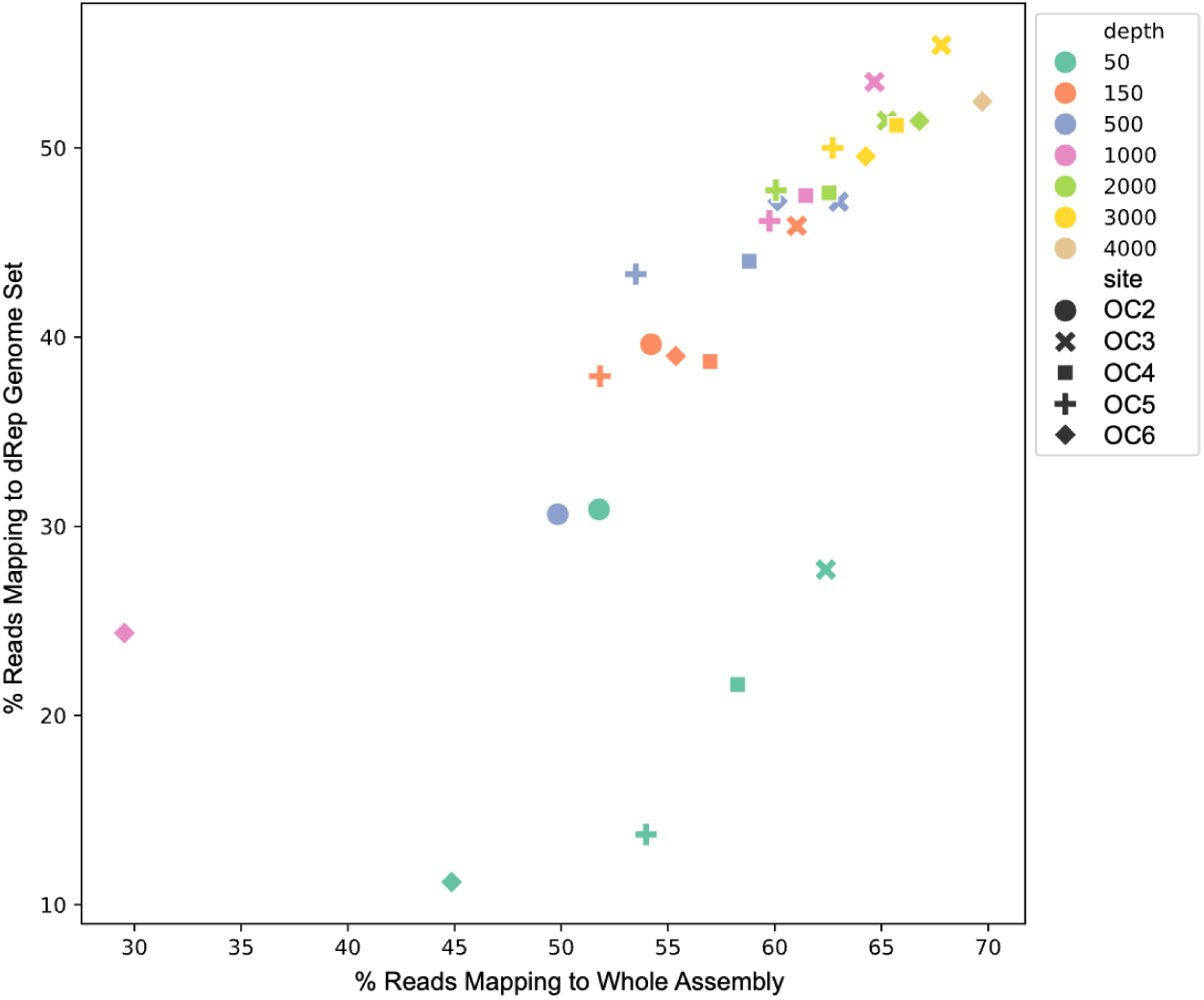
Percentage of trimmed sequencing reads recruited to the entire metagenomic assembly compared against percentage of trimmed sequencing reads recruited by the dereplicated genome set of 1195 genomes that are >50% complete and <25% contaminated. Samples are color coded by depth, and the different shapes represent the different sites from across the transect.

**Supplementary Figure S2:**
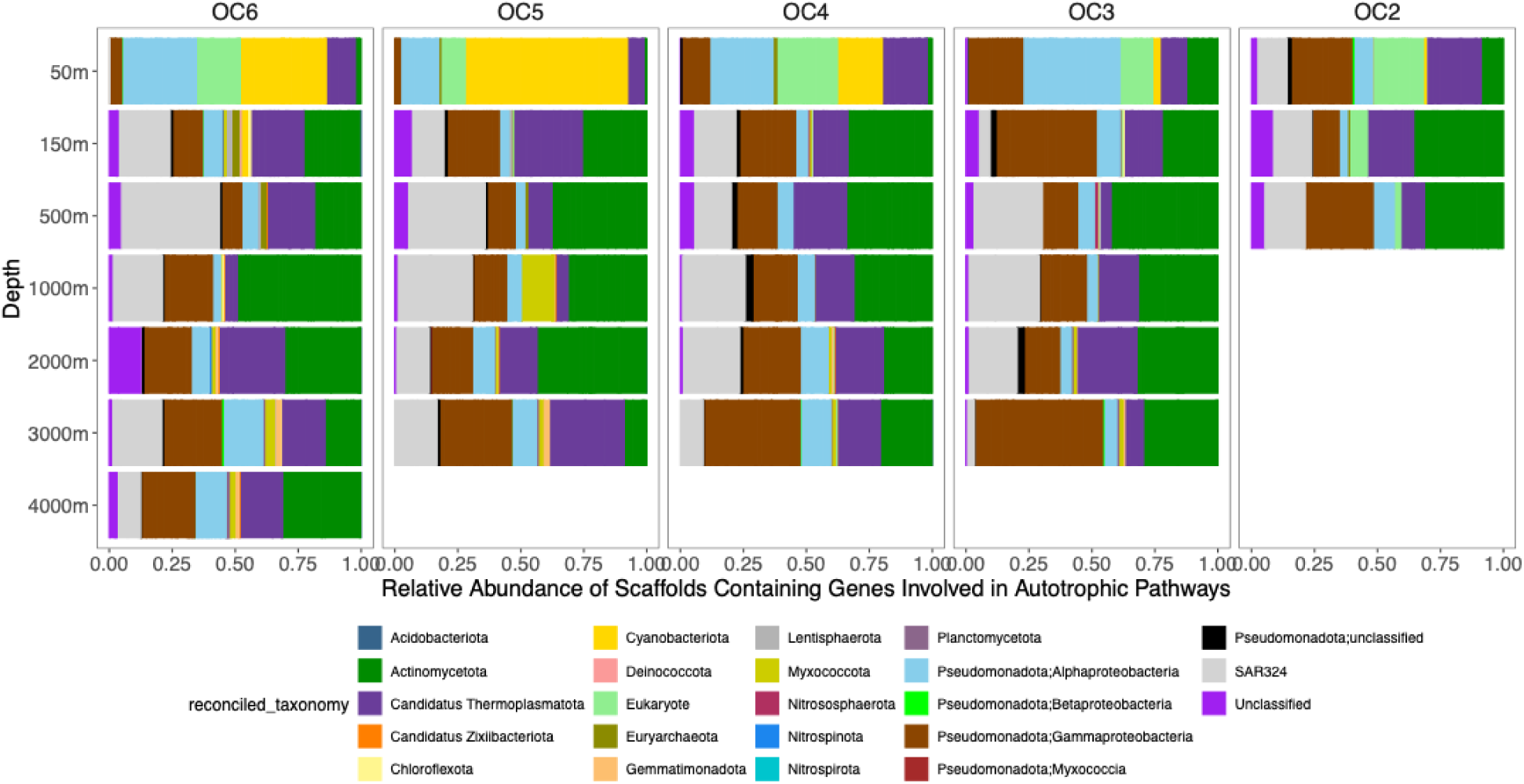
Presence and taxonomic identity of scaffolds encoding genes involved in autotrophic pathways across the transect, displayed as the abundance of any given scaffold relative to the coverage of all scaffolds that encode an autotrophy associated gene. Reconciled taxonomy is displayed at the phylum level as specified by NCBI, with the exception of the Pseudomadota (aka Proteobacteria), which are classified at the class level.

**Supplementary Figure S3:**
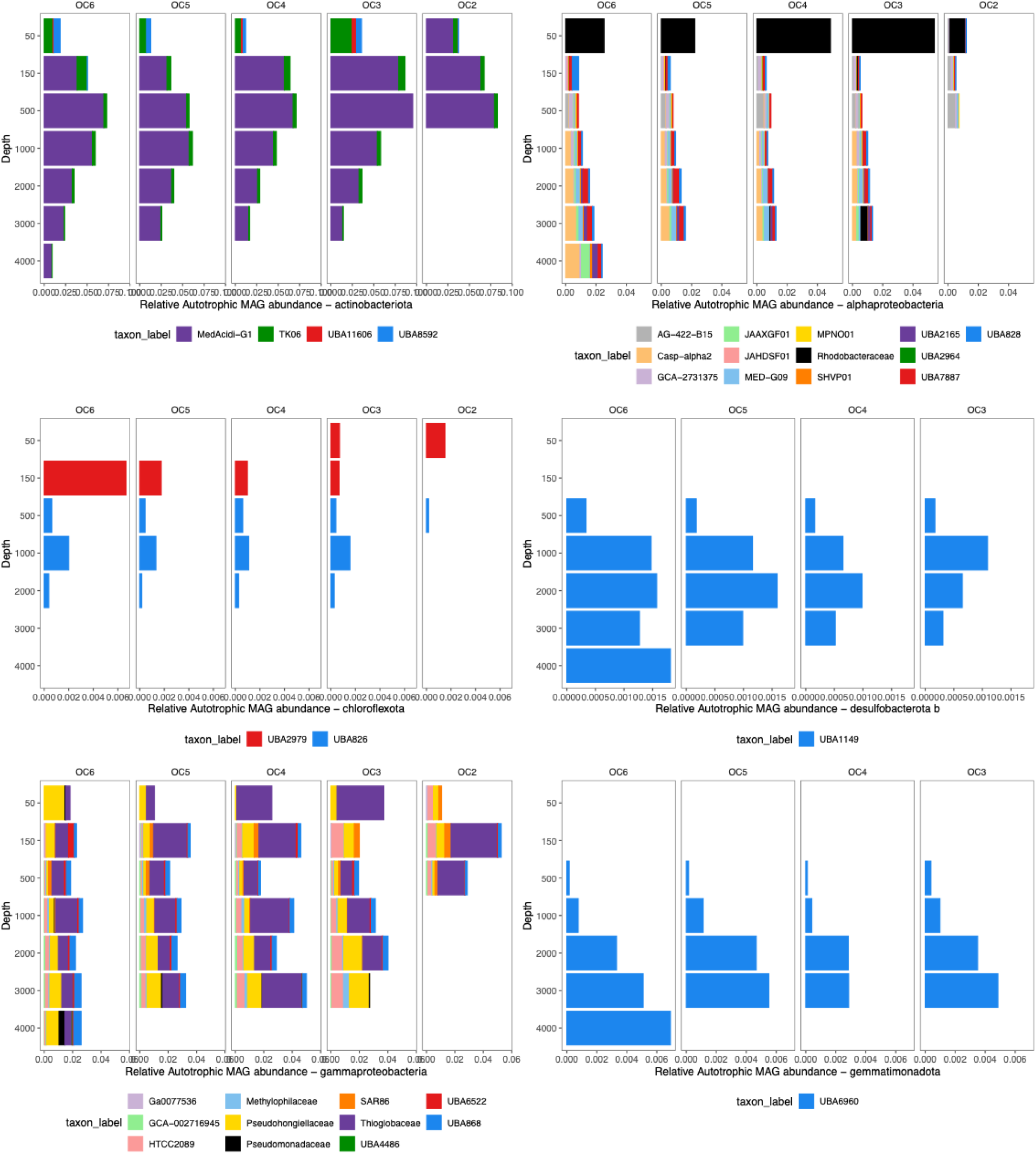
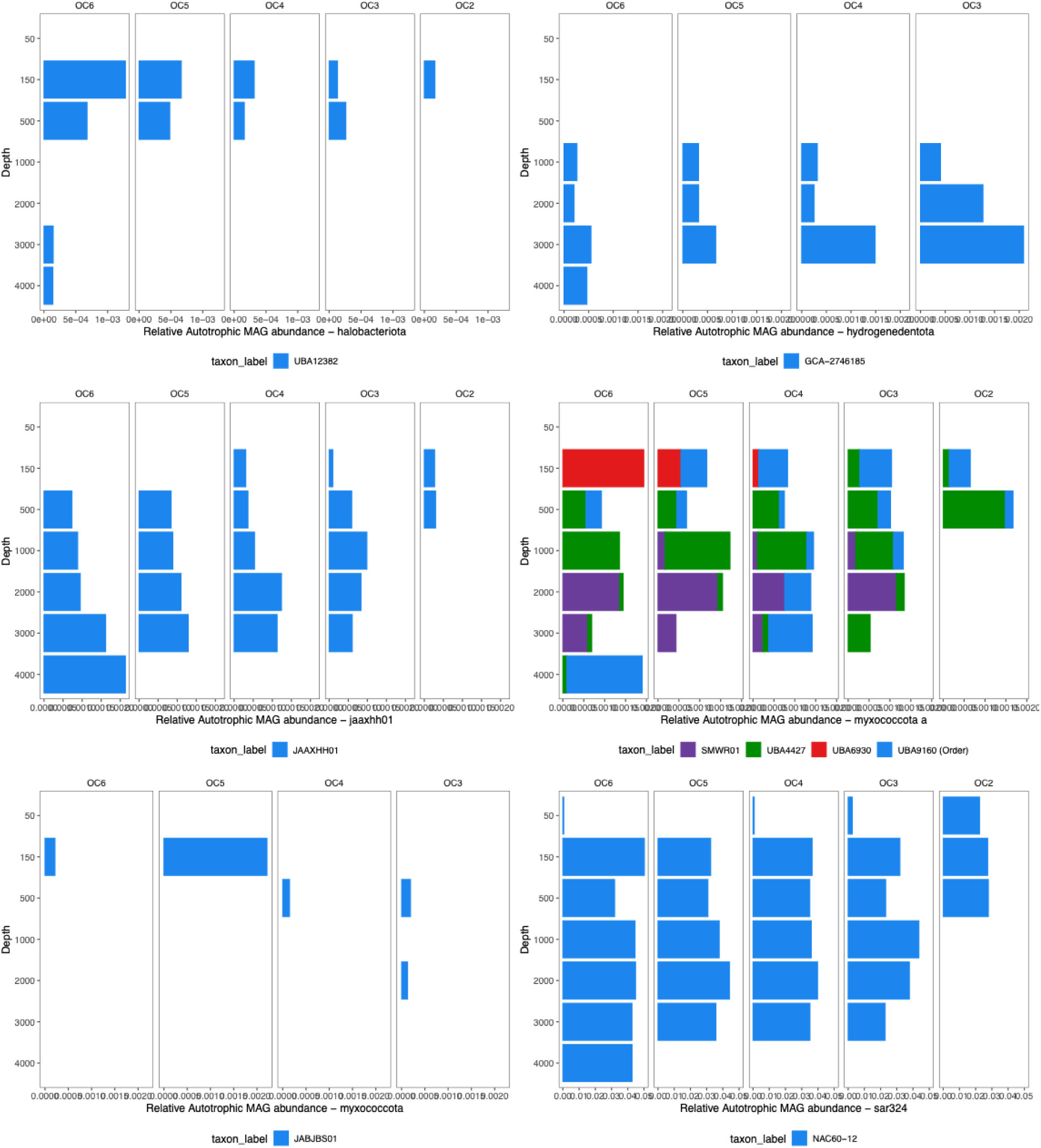
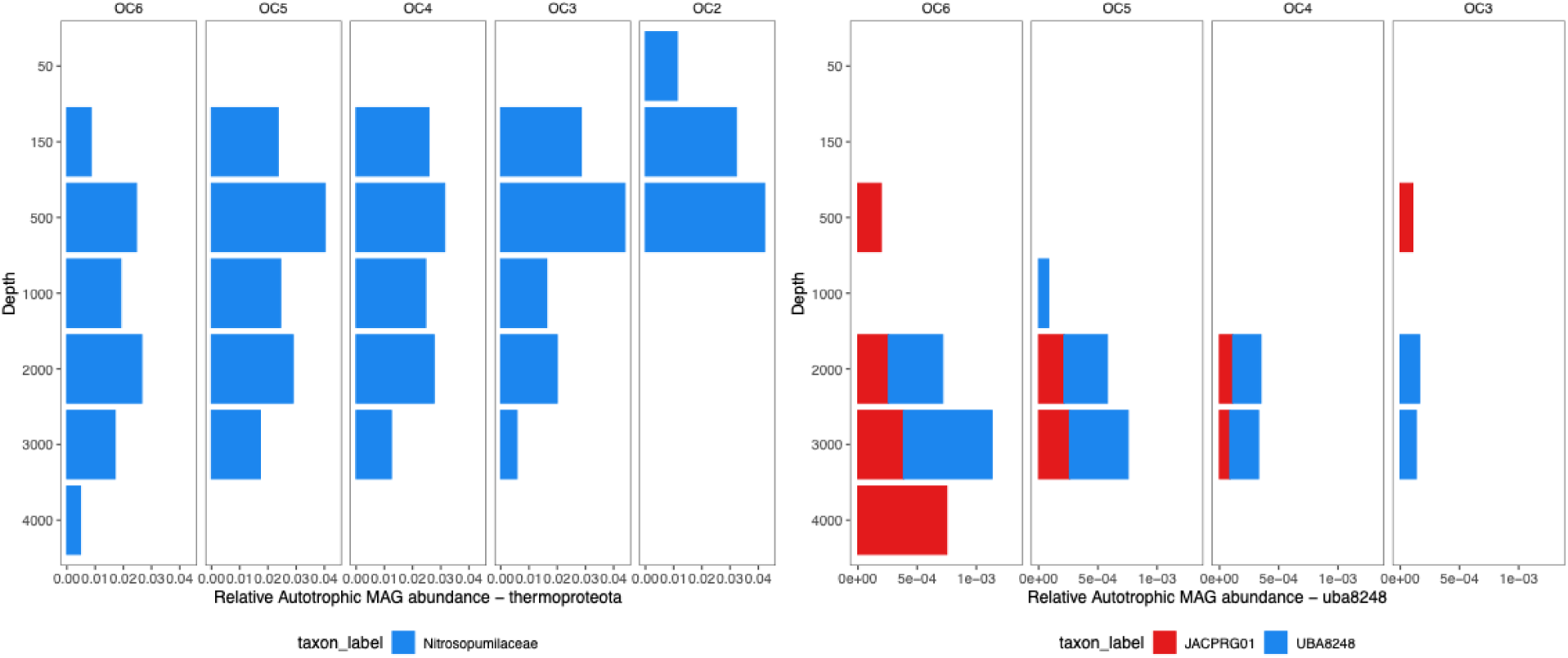
Relative abundance of putatively chemoautotrophic genomes across the transect relative to the total community. One plot per taxonomic phyla (except the Proteobacteria, which is broken up into the Alpha- and Gamma-proteobacteria), with MAGs color coded by family (unless Family was unclassified, in which case it’s plotted by order). Relative abundance is presented as a fraction (ranging from 0-1), not a percentage.

**Supplementary Figure S4:**
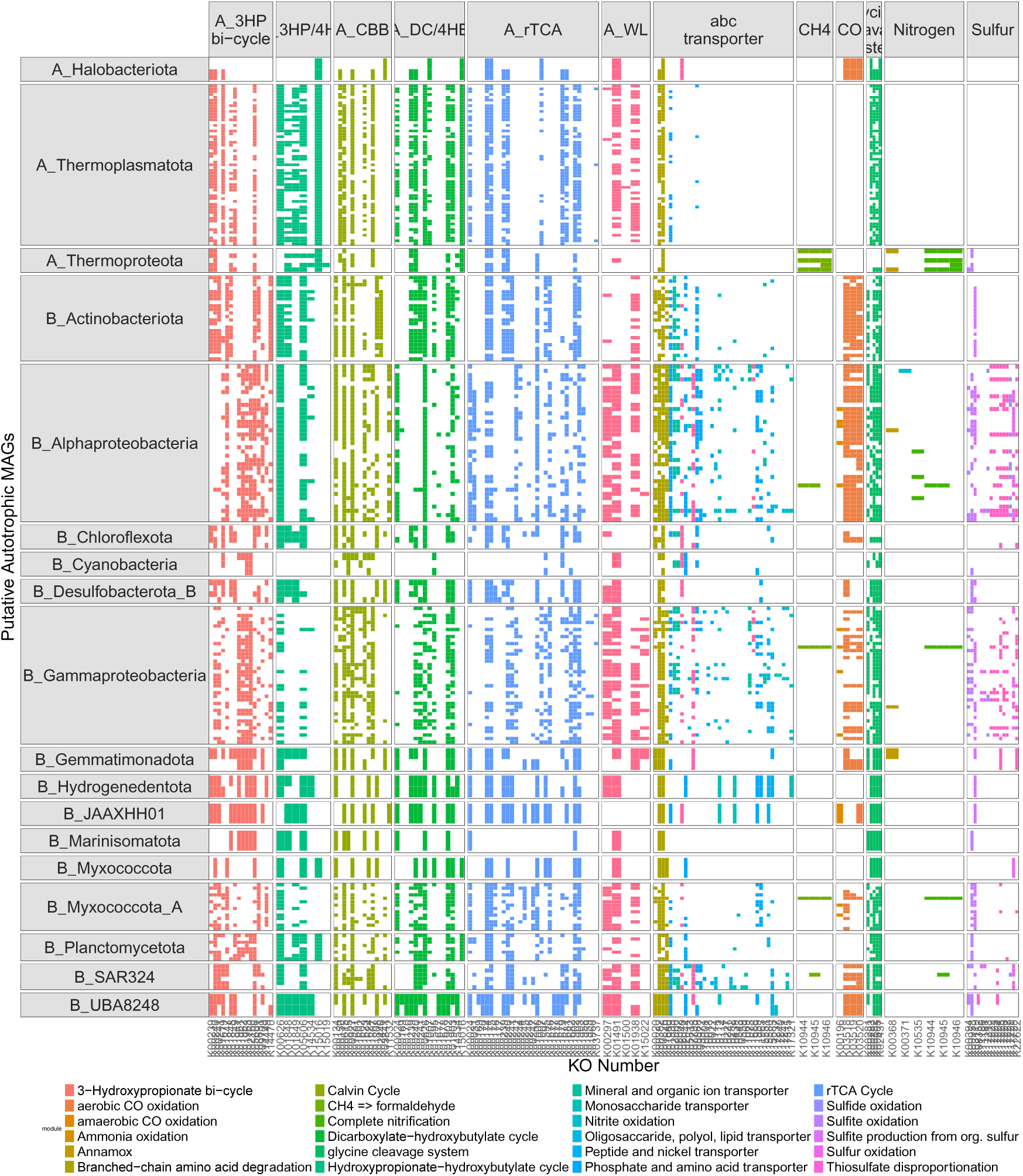
Metabolic potential of 212 MAGs encoding a marker gene for a DIC fixation pathway (similar to main text Figure 4), expanded to include the entirety of ICF pathways, not just marker genes, as well as genes associated with organic matter specific ABC transporters and the glycine cleavage system. Each row represents a single MAG, grouped by taxonomic lineage. Each column represents a single KO associated with a given catabolism or pathway of interest, grouped into larger metabolic functions. Boxes are colored coded by KEGG module aka metabolic pathway.

**Supplementary Figure S5:**
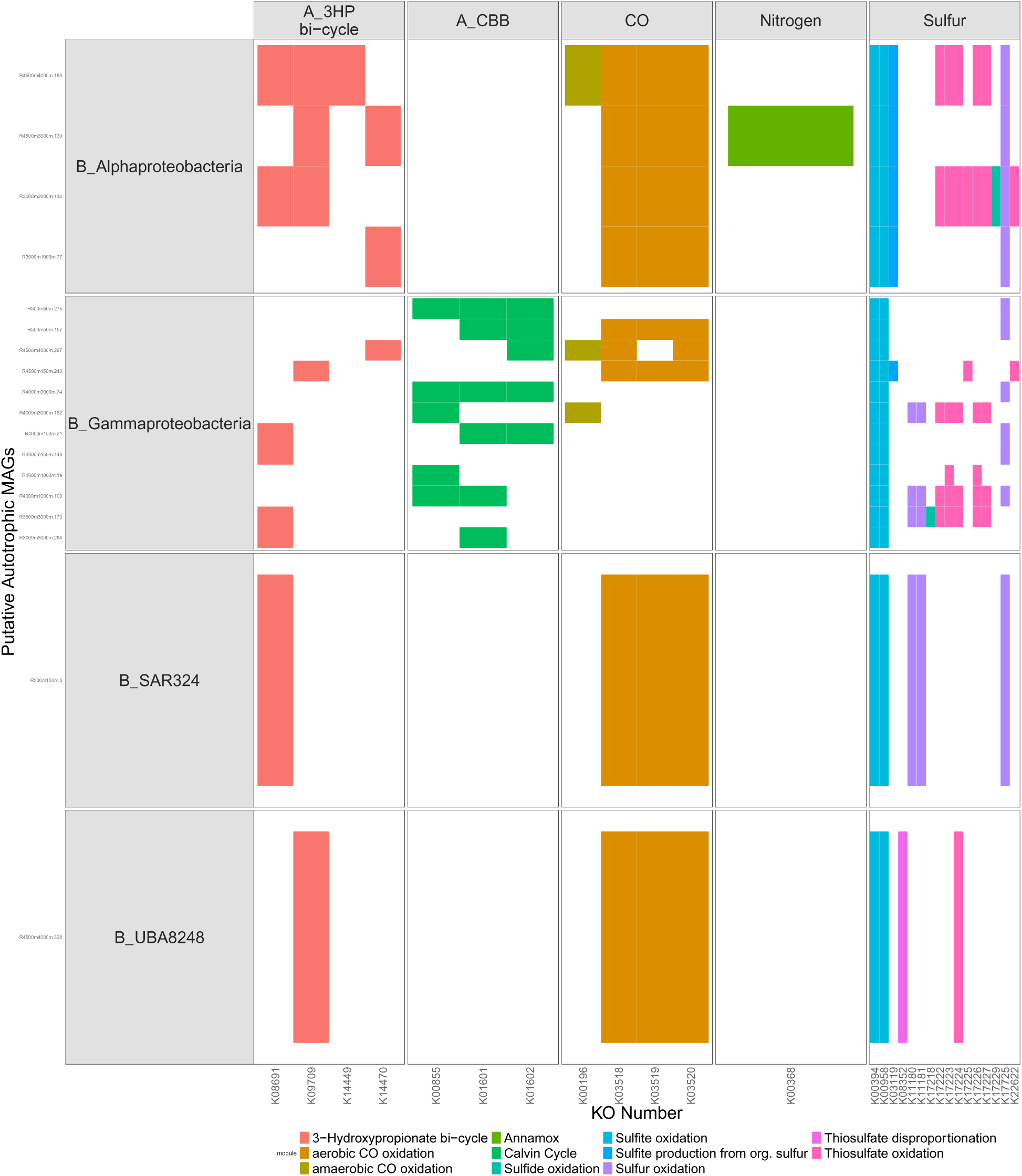
Metabolic potential of 18 putatively chemoautotrophic MAGs (similar to main text Figure 4), restricted to those determined to be capable of sulfite oxidation due to the presence of both *aprA* and *sat*. Each row represents a single MAG, grouped by taxonomic lineage. Each column represents a single KO associated with a given catabolism or pathway of interest, grouped into larger metabolic functions. Boxes are colored coded by KEGG module aka metabolic pathway.

